# Empirical Persistence Thresholds in Urban Arbovirus Dynamics: The Interplay of Population Size, Climate, and Urban Hierarchy in Brazil

**DOI:** 10.64898/2026.05.05.722903

**Authors:** César Castilho, João Gondim

**Affiliations:** Departmento de Matemática,Universidade federal de Pernambuco (UFPE), Cidade Universitária, Recife, Pernambuco, Brazil, 50670-420

**Keywords:** arboviruses, dengue, chikungunya, zika, epidemiological persistence, critical community size, urban networks, metapopulation dynamics, climate variability, Brazil, Wasserstein distance, persistence thresholds

## Abstract

The classical concept of Critical Community Size (CCS) as formulated by Bartlett defines the minimum host population required for a pathogen to persist endemically without stochastic extinction. While this framework successfully described directly transmitted childhood infections in relatively isolated populations, it is increasingly inadequate for modern urban systems characterized by strong connectivity between cities. Pathogens circulating in highly connected urban networks can repeatedly re-emerge through spatial reintroduction even when local transmission temporarily fades out. In such systems, persistence is inherently probabilistic and influenced simultaneously by population size, environmental suitability, and network connectivity. In this study, we develop a generalization of the CCS concept, the Empirical Persistence Threshold (EPT), and apply it to three of the main arboviruses circulating in Brazil—dengue, chikungunya, and Zika—over the period 2017–2024. The Empirical Persistence Threshold generalizes the classical notion of critical community size by replacing a single deterministic threshold with a probabilistic, datadriven measure. Instead of asking for the minimum population at which persistence is guaranteed, EPT characterizes the lower tail of the population distribution among municipalities that empirically sustain transmission. Using weekly incidence data from thousands of municipalities, we transform temporal incidence series into binary sequences describing the presence or absence of reported transmission. For each municipality, we characterize persistence through the empirical distribution of run lengths of consecutive weeks with reported cases. Distances between run-length distributions are computed using the Wasserstein-1 metric, allowing a geometrically meaningful comparison between persistence profiles, and municipalities are grouped into epidemiological regimes using hierarchical clustering methods. Across all three arboviruses, we identify two robust regimes: one exhibiting sporadic and recurrent epidemic transmission, and the other exhibiting sustained persistent transmission. We then estimate the population scales associated with each persistence regime. The analysis is further extended to evaluate how persistence thresholds vary across climate regimes (Köppen classification) and urban hierarchy levels (REGIC). This framework allows the estimation of probabilistic persistence thresholds analogous to CCS, but adapted to connected urban systems. We define the Empirical Persistence Threshold as lower quantiles of the population distribution among municipalities in the persistent regime, and additionally estimate persistence thresholds based on regime membership probabilities. Results reveal strong interactions between population size, climate, and urban connectivity. Dengue exhibits the lowest persistence thresholds, Zika intermediate thresholds, and chikungunya the highest thresholds. These findings demonstrate that pathogen persistence in modern urban systems cannot be described by a single deterministic population threshold. Instead, persistence emerges from the joint effects of demographic scale, environmental suitability, and network position within metapopulation systems.

**Author Summary:** Infectious diseases often require a minimum population size to persist locally, a concept known as the critical community size (CCS). This idea was developed for relatively isolated populations, but modern cities form highly connected networks where diseases can repeatedly reappear even after local transmission disappears.

In this study, we introduce the Empirical Persistence Threshold (EPT), a data-driven approach that replaces the idea of a single fixed threshold with a probabilistic description of persistence. Instead of focusing on case counts, we analyze how long transmission persists over time in each municipality.

Using weekly data for dengue, chikungunya, and Zika across Brazil from 2017 to 2024, we identify distinct patterns of transmission persistence and estimate the population levels associated with sustained transmission. We also examine how these thresholds vary with climate and urban structure.

Our results show that persistence depends not only on population size, but also on environmental conditions and the position of cities within the urban network.

## 1 Introduction

Understanding the mechanisms that allow infectious diseases to persist in host populations is a central problem in epidemiology. A classical framework addressing this question is the critical community size (CCS), defined as the minimum population required to sustain endemic transmission without stochastic extinction [1]. Originally developed for directly transmitted infections in relatively isolated populations, CCS successfully explained the contrast between persistent transmission in large cities and recurrent fadeouts in smaller communities.

However, the assumptions underlying CCS are difficult to reconcile with modern urban systems. Contemporary cities are embedded in highly connected mobility networks, where frequent human movement leads to rapid spatial reintroductions of pathogens [2]. In such settings, local extinction does not imply epidemiological independence, and persistence emerges from the interaction between local transmission dynamics and spatial coupling. These limitations are particularly relevant for arboviral diseases such as dengue, chikungunya, and Zika, whose transmission depends not only on host population size but also on vector ecology, climate variability, and urban connectivity [4, 5].

As a result, the notion of a single deterministic population threshold separating persistence from extinction becomes inadequate. Instead, persistence should be interpreted as a probabilistic and multidimensional phenomenon shaped by demographic, environmental, and network factors [3].

In this study, we propose a data-driven framework to characterize epidemiological persistence in large, interconnected urban systems. Our approach is based on the temporal structure of transmission, rather than on incidence magnitude or binary endemic classifications. Weekly incidence time series are transformed into binary presence– absence sequences, from which we extract the distribution of consecutive weeks with reported cases (run-length distributions). These distributions provide a compact yet informative representation of persistence, capturing the duration and continuity of transmission.

To compare municipalities, we employ the Wasserstein-1 distance from optimal transport theory [6], which allows for a geometrically meaningful comparison between run-length distributions. This choice is crucial: unlike traditional metrics, the Wasserstein distance preserves the notion of temporal proximity between run lengths, enabling a more faithful comparison of persistence patterns. Using these distances, we identify distinct epidemiological regimes through hierarchical clustering [7] and characterize their structure via non-metric multidimensional scaling (NMDS) [8].

Building on this empirical characterization, we introduce the concept of the Empirical Persistence Threshold (EPT). Instead of a single deterministic threshold, EPT is defined from the observed distribution of population sizes among municipalities exhibiting sustained transmission. Lower quantiles of this distribution (e.g., 1%, 5%, and 10%) provide robust estimates of the minimum population scales associated with persistence, explicitly acknowledging its probabilistic nature.

We apply this framework to dengue, chikungunya, and Zika across Brazilian municipalities from 2017 to 2024, integrating demographic, climatic (Köppen–Geiger classification [9]), and urban hierarchy (REGIC) variables [10].

Our results reveal that persistence thresholds vary systematically across diseases and depend strongly on the interplay between population size, climate, and urban structure.

Taken together, these findings challenge the classical CCS paradigm and support a conceptual shift toward viewing persistence as an emergent property of complex, interconnected systems. The run-length distributions, combined with optimal transport metrics, provide a novel and generalizable methodology for quantifying persistence in modern epidemiological landscapes.

## 2 Methods

We analyzed weekly notified cases of dengue, chikungunya, and Zika for Brazilian municipalities from 2017 to 2024, obtained from the Brazilian national surveillance system (SINAN) [11]. To ensure temporal consistency, only municipalities present in all years were retained, yielding an intersection dataset for each disease. For each municipality, we assembled the population size, Köppen climate classification and urban hierarchy classification based on REGIC [10]. The REGIC (Regi∼oes de Influ^encia das Cidades) classification describes the hierarchical organization of Brazilian cities according to their economic and functional influence within the national urban network. Categories include metropolitan centers, regional capitals, sub-regional centers, and local centers. Together, these variables capture three complementary dimensions that are expected to influence pathogen persistence: demographic scale, environmental suitability and connectivity within the urban system. These covariates are not directly used in the construction of the persistence regimes, which are defined solely from temporal patterns of incidence via run-length distributions and Wassersteinbased clustering. Instead, they provide an external explanatory framework, allowing us to interpret the resulting regimes in terms of demographic, environmental, and urban connectivity factors in subsequent analyses.

### 2.1 Data and Run-length distributions

For each municipality *i* and week *t*, let *X*_*i*_ (*t*) denote the number of reported cases. We transformed each time series into a binary presence–absence sequence:

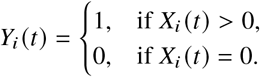

This transformation focuses the analysis on the temporal continuity of transmission rather than on incidence magnitude itself. Persistence was characterized through consecutive runs of weeks with reported cases.

A run of length *l* is defined as a maximal sequence of *l* consecutive weeks with *Y*_*i*_ (*t*) = 1. For each municipality, let *P*_*i*_ (*l*) denote the empirical distribution of run lengths:

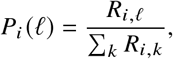

where *R*_*i*,*l*_ is the number of runs of length *l*. The distribution *P*_*i*_ (*l*) summarizes how transmission is temporally organized, distinguishing sporadic from sustained dynamics. Large metropolitan areas can produce extremely long runs, generating heavy tails in the distribution.

To mitigate this effect, we introduced a truncation level *L*_max_ defined as the smallest value for which the average cumulative mass of the run-length distributions across municipalities reaches 99%. This guarantees that most municipalities have nearly all their probability mass concentrated within 1, [*L*_max_], while only a small fraction display heavy-tailed behaviour beyond this range. That is, *L*_max_ is defined by

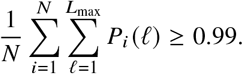

This retains 99% of the probability mass while removing extreme outliers.

The mean run length was also computed as

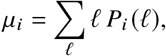

which represents the expected duration of a transmission episode.

### 2.2 Distance between persistence profiles

To compare municipalities, we used the one-dimensional Wasserstein-1 distance between run-length distributions. For municipalities *i* and *j*, this distance is given by

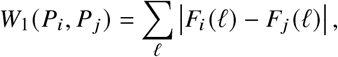

where *F*_*i*_ and *F*_*j*_ are the cumulative distributions of *P*_*i*_ and *P* _*i*_.

This metric preserves the ordering of run lengths, ensuring that differences between similar durations (e.g., 2 vs. 3 weeks) are treated as smaller than differences between distant durations (e.g., 2 vs. 20 weeks). This property makes it particularly suitable for comparing temporal persistence patterns.

### 2.3 Identification of persistence regimes

Pairwise Wasserstein distances were assembled into a dissimilarity matrix and used for hierarchical clustering with Ward’s minimum variance method (Ward.D2). This procedure groups municipalities with similar temporal persistence profiles. Cluster quality was evaluated using the silhouette coefficient. Across datasets, the most robust and interpretable solution corresponds to *k* = 2. The resulting clusters were interpreted as epidemiological regimes. In practice, two dominant regimes emerged:

- low persistence: characterized by short and fragmented runs;
- high persistence: characterized by longer and more continuous transmission.

### 2.4 Non-metric multidimensional scaling

To visualize the structure of persistence patterns, we applied non-metric multidimensional scaling (NMDS) to the Wasserstein dissimilarity matrix. NMDS provides a low-dimensional representation that preserves the rank order of dissimilarities, allowing qualitative assessment of regime separation and gradients of persistence.

### 2.5 Empirical Persistence Threshold (EPT)

To quantify the population scale associated with sustained transmission, we defined the Empirical Persistence Threshold (EPT).

Let *H* denote the set of municipalities classified in the high-persistence regime, and let *N*_*i*_ be the population of municipality *i*. The EPT is defined through lower quantiles of the population distribution within *H*:

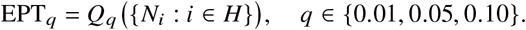

These quantiles provide complementary information:

- *q* = 0.01: extreme lower bound;
- *q* = 0.05: robust empirical threshold;
- *q* = 0.10: onset of consistent persistence.

This formulation replaces the notion of a single deterministic threshold by a probabilistic description of persistence.

### 2.6 Stratified analysis

To investigate how persistence depends on environmental and urban structure, we considered climate and urban hierarchy as explanatory variables.

Climate was represented using a collapsed Köppen classification, while urban structure was described using a collapsed REGIC hierarchy. These aggregations were designed to group categories with similar epidemiological and structural characteristics, improving interpretability and statistical robustness.

The resulting classifications are summarized in Table 1.

**Table 1:** Collapsed classifications used in the analysis. Urban hierarchy (REGIC) and climate (Köppen) categories were aggregated into broader groups to improve interpretability and statistical robustness.

| REGIC (Urban Hierarchy) |  | Köppen (Climate) |  |
| --- | --- | --- | --- |
| Level | Original Categories | Class | Original Classes |
| 1 | Metropolises (1A, 1B) | Equatorial | Af, Am |
| 2 | Regional Capitals (2A, 2B, 2C) | Tropical | Aw, As |
| 3 | Sub-regional Centers (3A, 3B) | Semi-arid | BSh |
| 4 | Zone Centers (4A, 4B) | Temperate | Cfa, Cfb, Cwa, Cwb |
| 5 | Local Centers (5) |  |  |

**Table 2:**
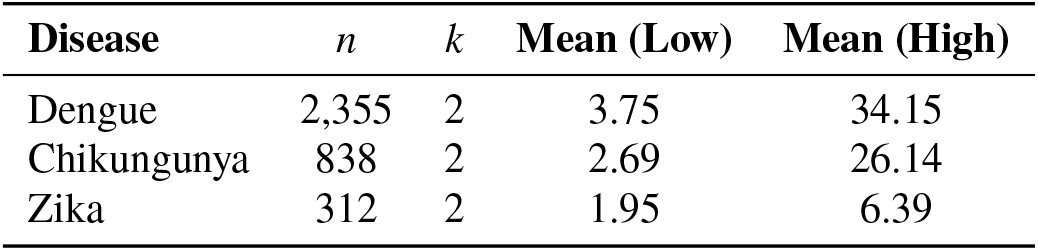
Comparison of persistence regimes across arboviruses. Values correspond to the mean run length (weeks) and cluster sizes obtained from hierarchical clustering.

**Table 3:** Cluster sizes for persistence regimes. The high-persistence regime is consistently rare across diseases.

| Disease | Low Persistence | High Persistence | Proportion High (%) |
| --- | --- | --- | --- |
| Dengue | 2,329 | 27 | 1.15 |
| Chikungunya | 828 | 10 | 1.19 |
| Zika | 293 | 19 | 6.09 |

To assess the role of environmental and urban factors, EPT was computed within strata defined by climate and urban hierarchy.

Climate was represented by a collapsed Köppen classification (Equatorial, Tropical, Semi-arid, Temperate), while urban structure was represented by a collapsed REGIC hierarchy (levels 1 to 5).

For each disease and each stratum *g*, EPT was estimated from the subset of municipalities in the high-persistence regime belonging to *g*, allowing evaluation of how persistence thresholds vary across climatic and urban contexts.

### 2.7 Interpretive framework

The overall analytical pipeline can therefore be summarized as follows (see Figure 1):

**Figure 1:**
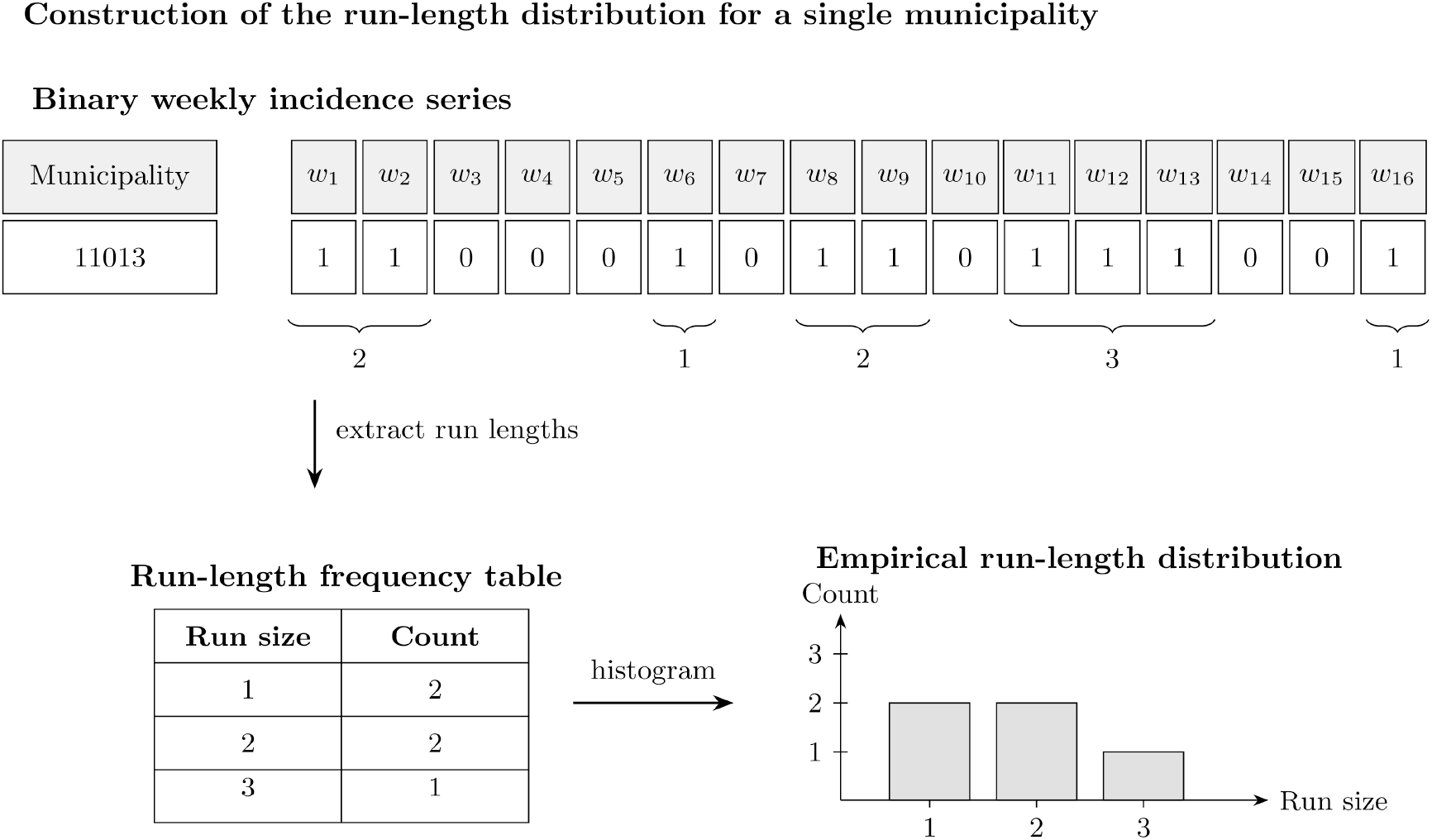
Construction of the run-length distribution from a binary weekly transmission sequence for a single municipality. Weekly incidence is converted into a binary presence–absence sequence, where consecutive weeks with reported cases define runs. These run lengths are extracted, tabulated, and transformed into an empirical distribution, providing a compact summary of the temporal persistence structure of local transmission.

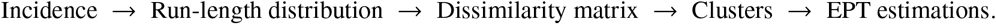

Starting from weekly incidence data, we transform the time series into a binary sequence indicating the presence or absence of reported transmission. Consecutive weeks with reported cases form runs, whose lengths capture the temporal continuity of transmission. The empirical distribution of these run lengths provides a compact representation of persistence.

As shown in Figure 2, municipalities in the persistent regime are characterized by distributions with substantial mass at longer run lengths, reflecting sustained transmission over extended periods. In contrast, low-persistence municipalities exhibit distributions concentrated at short run lengths, corresponding to sporadic and fragmented transmission. This representation forms the basis for the distance-based comparisons and clustering analyses developed in this study.

**Figure 2:**
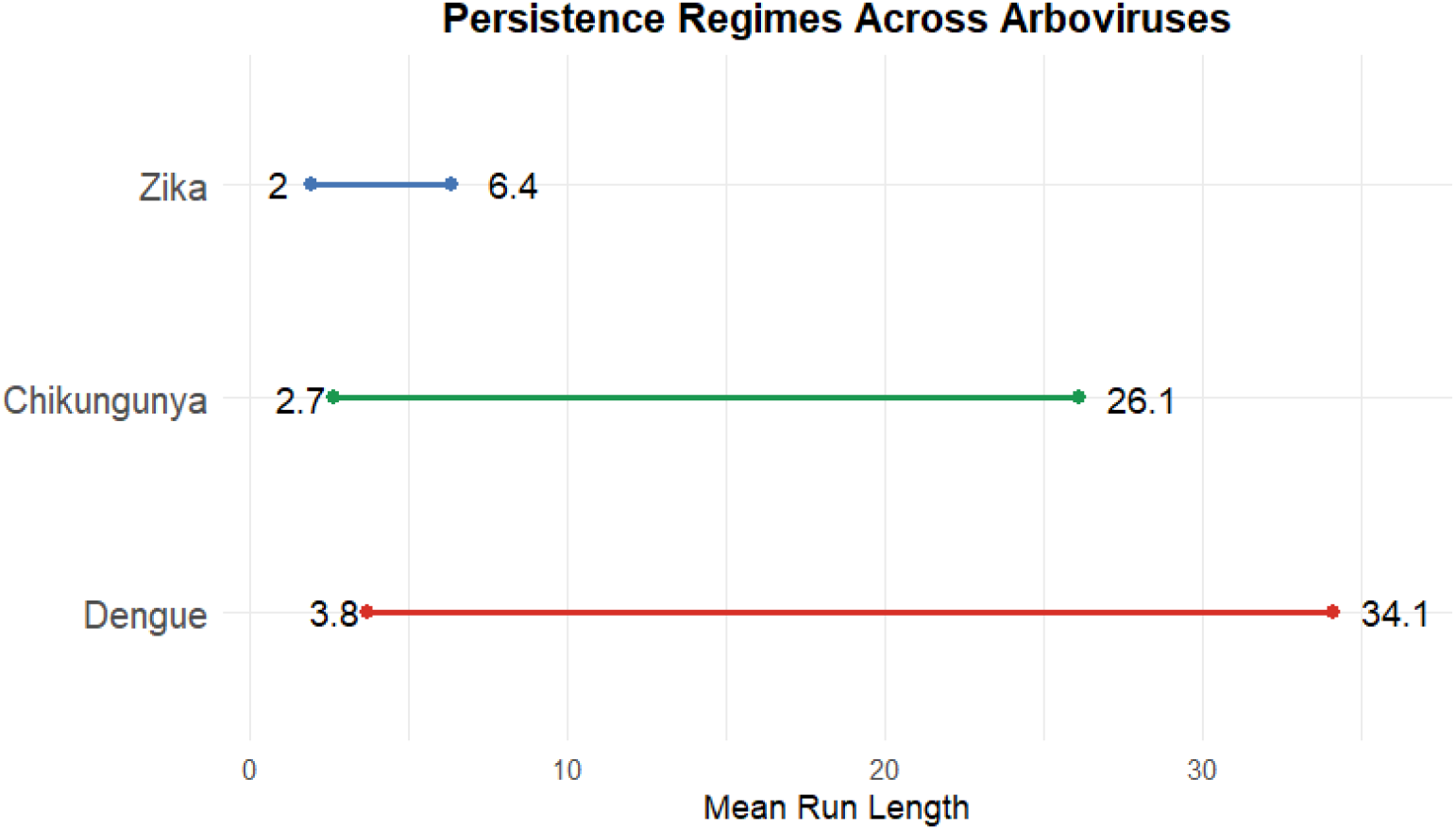
Run-length values for dengue, zika and chykungunya.

Additional methodological details and robustness analyses are provided in the Supplementary Material: S1 Text. This framework combines a temporally explicit representation of transmission persistence with a distribution-sensitive metric. This allows for an empirically grounded threshold concept. It replaces the idea of a universal critical community size by a more realistic description of persistence as a multidimensional and context-dependent phenomenon.

## 3 Results

### 3.1 Global structure of persistence patterns

The intersection dataset comprised 2,355 municipalities for dengue, 838 for chikungunya, and 312 for Zika. These differences reflect both the spatial extent of transmission and the temporal consistency of reported cases across the study period (2017–2024).

The run-length distributions revealed substantial heterogeneity in the temporal structure of transmission across Brazilian municipalities for all three arboviruses as shown in Figure 2 shows. While some municipalities exhibited predominantly short runs of presence, indicating sporadic or fragmented transmission, others displayed substantial probability mass at longer run lengths, consistent with sustained epidemiological activity.

This heterogeneity was already apparent at the level of summary statistics. The mean run length µ_*i*_ varied widely across municipalities, spanning from values close to one week—indicative of isolated transmission events—to several tens of weeks, reflecting prolonged periods of continuous transmission. This broad spectrum of behaviors suggests that persistence is not a binary property but rather a continuous feature of the underlying dynamics.

### 3.2 Identification of persistence regimes

Hierarchical clustering based on Wasserstein distances between run-length distributions consistently revealed a clear separation between two dominant persistence regimes across all diseases.

The first regime, hereafter referred to as the low-persistence regime, was characterized by distributions concentrated at small run lengths, typically dominated by runs of one to three weeks. Municipalities in this regime exhibited intermittent transmission, with frequent interruptions and recurrent fade-outs.

The second regime, the high-persistence regime, displayed a markedly different structure, with substantial probability mass extending toward longer run lengths. These municipalities experienced sustained transmission episodes lasting several consecutive weeks or months.

Despite the simplicity of this two-regime classification, it captured the dominant structure of the data and provided a robust basis for subsequent analyses.

Clustering of run-length distributions revealed a consistent two-regime structure across all three arboviruses, comprising a dominant low-persistence regime and a rare high-persistence regime (Table 4).

**Table 4:**
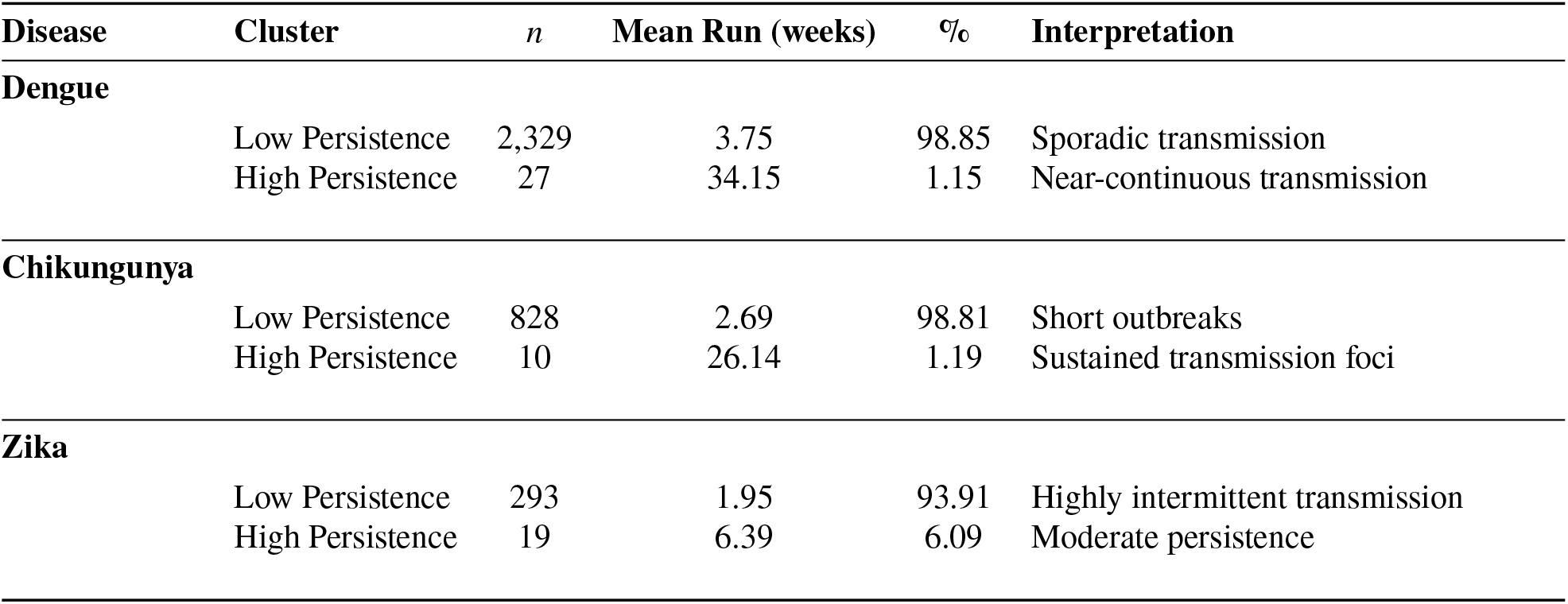
Run-length statistics and cluster structure for persistence regimes across arboviruses. Each disease exhibits two regimes: low persistence and high persistence.

Despite this shared qualitative structure, the quantitative characteristics of persistence differed markedly between diseases. Dengue exhibited the strongest persistence signal. While the majority of municipalities (98.85%) belonged to the low-persistence regime, characterized by short mean runs of approximately 3.75 weeks, a small subset (1.15%) displayed extremely long runs, with a mean duration exceeding 34 weeks. These values indicate near-continuous transmission in a limited number of municipalities.

Chikungunya showed a similar but less pronounced pattern. The high-persistence regime comprised only 1.19% of municipalities, with a mean run length of approximately 26 weeks. Although substantial, this level of persistence is notably lower than that observed for dengue, suggesting a reduced capacity for sustained transmission.

In contrast, Zika exhibited a fundamentally different persistence profile. Although a two-regime structure was still identifiable, the high-persistence cluster accounted for a larger fraction of municipalities (6.09%) but with much shorter mean runs (approximately 6.39 weeks). This indicates that even in its most persistent settings, Zika transmission remains comparatively intermittent.

Overall, these results demonstrate that while all three arboviruses share a common structural organization into persistence regimes, the strength and epidemiological significance of these regimes vary substantially. Dengue emerges as the most structurally persistent disease, followed by chikungunya, whereas Zika displays a more intermittent persistence pattern.

### 3.3 Geometric structure of persistence (NMDS)

To assess the geometric structure of persistence regimes, we projected the Wasserstein distance matrices into a two-dimensional space using non-metric multidimensional scaling (NMDS). Two datasets were considered: the full intersection dataset, containing all municipalities consistently present across the study period, and the core dataset, obtained after removing extreme persistence profiles to evaluate the robustness of the clustering structure.

The NMDS representation of the Wasserstein distance matrix provided a low-dimensional visualization of persistence patterns and confirmed the existence of well-separated regimes.

In all three diseases, municipalities formed a structured continuum in the NMDS space, with a clear gradient from low to high persistence. The high-persistence municipalities occupied a distinct region of the space, separated from low-persistence municipalities along the primary NMDS axis.

Stress values were low, indicating that the two-dimensional representation preserved the rank structure of the original dissimilarities and provided a faithful summary of the persistence landscape.

As shown in Figure 3, the NMDS ordinations reveal a clear separation between persistence regimes.

**Figure 3:**
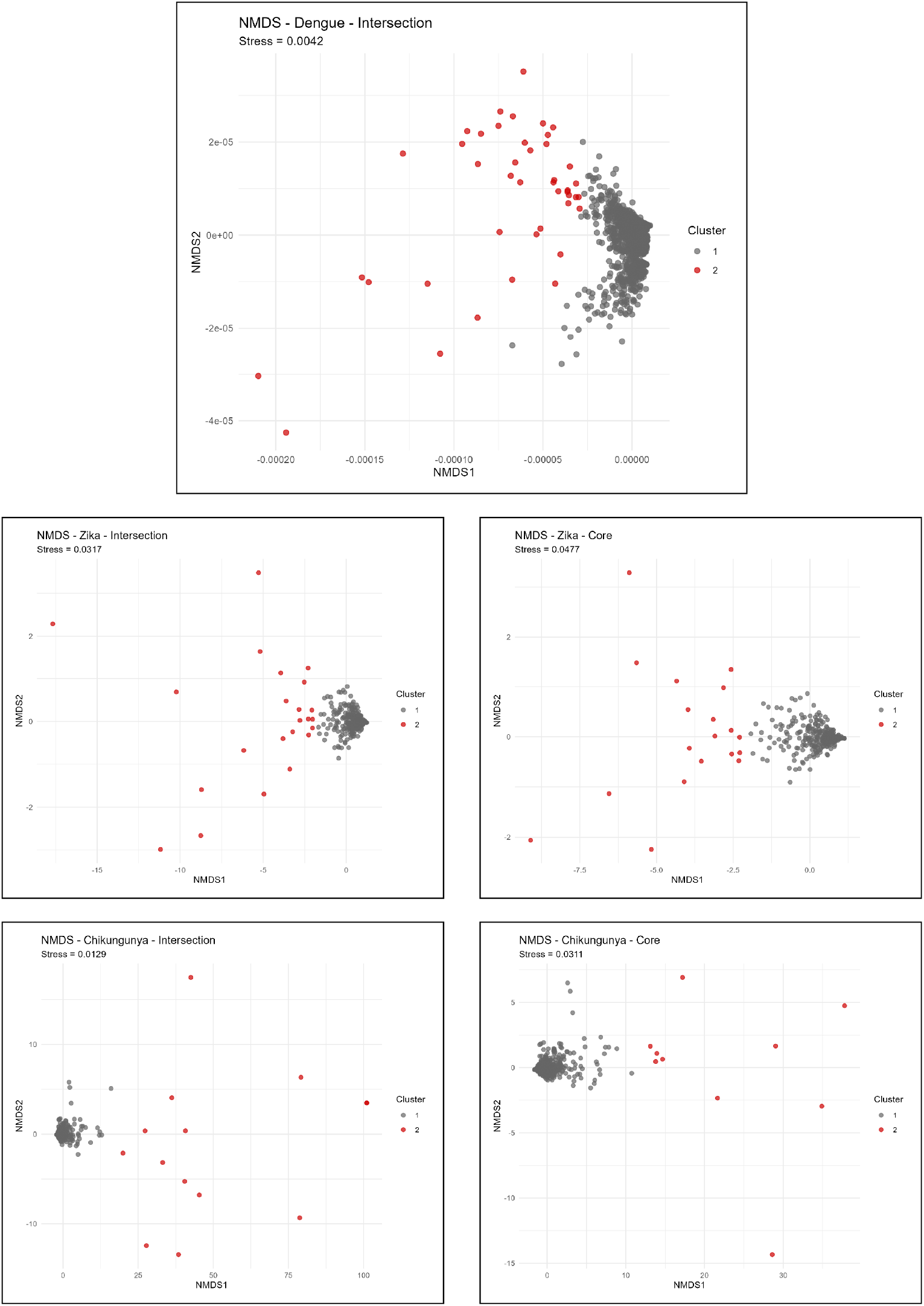
Non-metric multidimensional scaling (NMDS) representations of persistence regimes for dengue, Zika, and chikungunya based on Wasserstein distances between municipal run-length distributions. Each panel shows a two-dimensional ordination of municipalities colored according to persistence regime. Across all arboviruses, municipalities organize along a clear gradient from low to high persistence, with high-persistence municipalities forming a distinct and well-separated region of the space. The similarity between intersection and core datasets indicates that the clustering structure is robust to the removal of extreme persistence profiles.

For dengue, the NMDS projections for the intersection and core datasets are identical, reflecting the absence of extreme municipalities requiring removal. This indicates that the clustering structure is intrinsically robust. The NMDS configuration reveals a clear and well-separated two-regime structure, with a compact low-persistence cluster and a distinctly isolated high-persistence group.

In chikungunya and Zika the core dataset differs slightly from the intersection dataset due to the removal of a small number of extreme municipalities.

### 3.4 Empirical Persistence Thresholds (EPT)

The Empirical Persistence Threshold (EPT) provided a quantitative characterization of the population scale associated with sustained transmission.

The results suggest that persistence in arboviral diseases should be understood in terms of regime structure rather than smooth probabilistic transitions. In highly connected urban systems, persistence is better described by threshold effects and regime boundaries than by classical CCS concepts.

This has important implications for epidemiological modeling and public health planning. In particular, interventions may need to be targeted at specific municipalities that act as persistence hubs, rather than relying on population-based thresholds alone.

For dengue, persistence was observed across a wide range of population sizes, including relatively small municipalities. The estimated thresholds (Table 5) indicate that dengue can sustain transmission at substantially lower population levels than the other arboviruses. These values suggest that dengue can persist in relatively small populations, particularly under favorable environmental conditions.

**Table 5:** Empirical persistence thresholds based on population size.

| Disease | Minimum | 1% | 5% | 10% |
| --- | --- | --- | --- | --- |
| Dengue | 15,356 | 24,894 | 63,044 | 110,732 |
| Zika | 4,203 | 20,274 | 84,560 | 97,830 |
| Chikungunya | 218,049 | 219,603 | 225,817 | 233,585 |

**Table 6:** EPT stratified by climate zones (collapsed Köppen classification).

| Group | Disease | n | Min | Q01 | Q05 | Q10 |
| --- | --- | --- | --- | --- | --- | --- |
| Equatorial | Chikungunya | 4 | 218,049 | 222,449 | 240,049 | 262,049 |
| Tropical | Chikungunya | 6 | 235,311 | 236,874 | 243,124 | 250,936 |
| Equatorial | Dengue | 7 | 98,916 | 114,890 | 178,784 | 258,653 |
| Temperate | Dengue | 3 | 108,708 | 121,075 | 170,545 | 232,382 |
| Tropical | Dengue | 17 | 9,132 | 10,128 | 14,111 | 19,569 |
| Equatorial | Zika | 10 | 98,916 | 99,602 | 102,346 | 105,777 |
| Semi-arid | Zika | 1 | 278,587 | 278,587 | 278,587 | 278,587 |
| Temperate | Zika | 1 | 1,190,496 | 1,190,496 | 1,190,496 | 1,190,496 |
| Tropical | Zika | 7 | 4,203 | 9,560 | 30,989 | 57,774 |

**Table 7:** EPT stratified by urban hierarchy (REGIC, collapsed levels).

| Disease | REGIC | n | Min | Q01 | Q05 | Q10 |
| --- | --- | --- | --- | --- | --- | --- |
| Dengue | 1 | 9 | 15,356 | 24,893.60 | 63,044.00 | 11,0732.0 |
| Dengue | 2 | 15 | 2,2378 | 31,029.16 | 65,633.80 | 93,986.4 |
| Dengue | 3 | 3 | 9,132 | 10,449.34 | 15,718.70 | 22,305.4 |
| Chikungunya | 1 | 6 | 218,049 | 225,382.40 | 254,716.00 | 291,383.0 |
| Chikungunya | 2 | 4 | 235,311 | 236,248.50 | 239,998.50 | 244,686.0 |
| Zika | 1 | 7 | 106,539 | 131,153.88 | 229,613.40 | 352,687.8 |
| Zika | 2 | 10 | 4,203 | 20,296.89 | 84,672.45 | 165,141.9 |
| Zika | 3 | 2 | 93,488 | 93,542.28 | 93,759.40 | 94,030.8 |

In contrast, chikungunya exhibited substantially higher persistence thresholds (Table 5), suggesting that sustained transmission is largely restricted to larger host populations, consistent with more constrained persistence dynamics. Zika displayed intermediate behavior, with persistence thresholds between those of dengue and chikungunya (Table 5). These results demonstrate that persistence thresholds are not universal across arboviruses but instead reflect disease-specific transmission dynamics.

The empirical persistence thresholds (EPT) as a function of population size for the three arboviruses is summarized in Figure 4. The thresholds are reported as lower quantiles, capturing the lower envelope of population sizes associated with sustained transmission.

**Figure 4:**
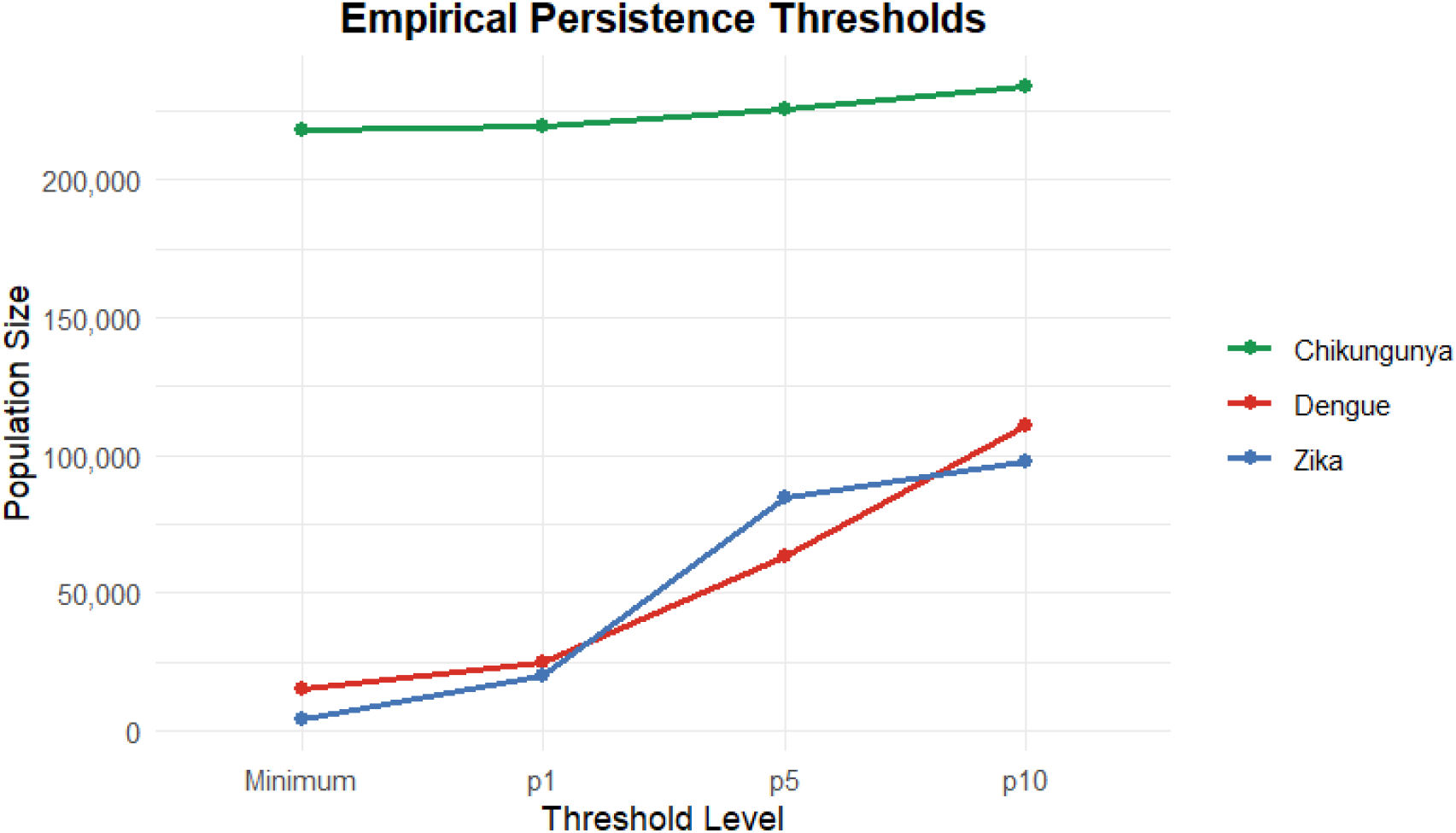
Empirical persistence thresholds (EPT) as a function of population size for dengue, Zika, and chikungunya. Thresholds are shown at different quantile levels (minimum, 1%, 5%, and 10%). Chikungunya exhibits consistently higher thresholds, while dengue and Zika show lower and more gradual transitions.

### 3.5 Influence of climate on persistence

Stratification by Köppen climate classes revealed systematic variation in persistence thresholds across environmental conditions.

The variation of empirical persistence thresholds across climate zones is shown in Figure 5. The relationship is strongly disease-dependent. Zika exhibits pronounced sensitivity to climate, with substantially higher thresholds in temperate regions. In contrast, dengue and chikungunya display more moderate variation, suggesting that climate influences persistence differently across arboviruses.

**Figure 5:**
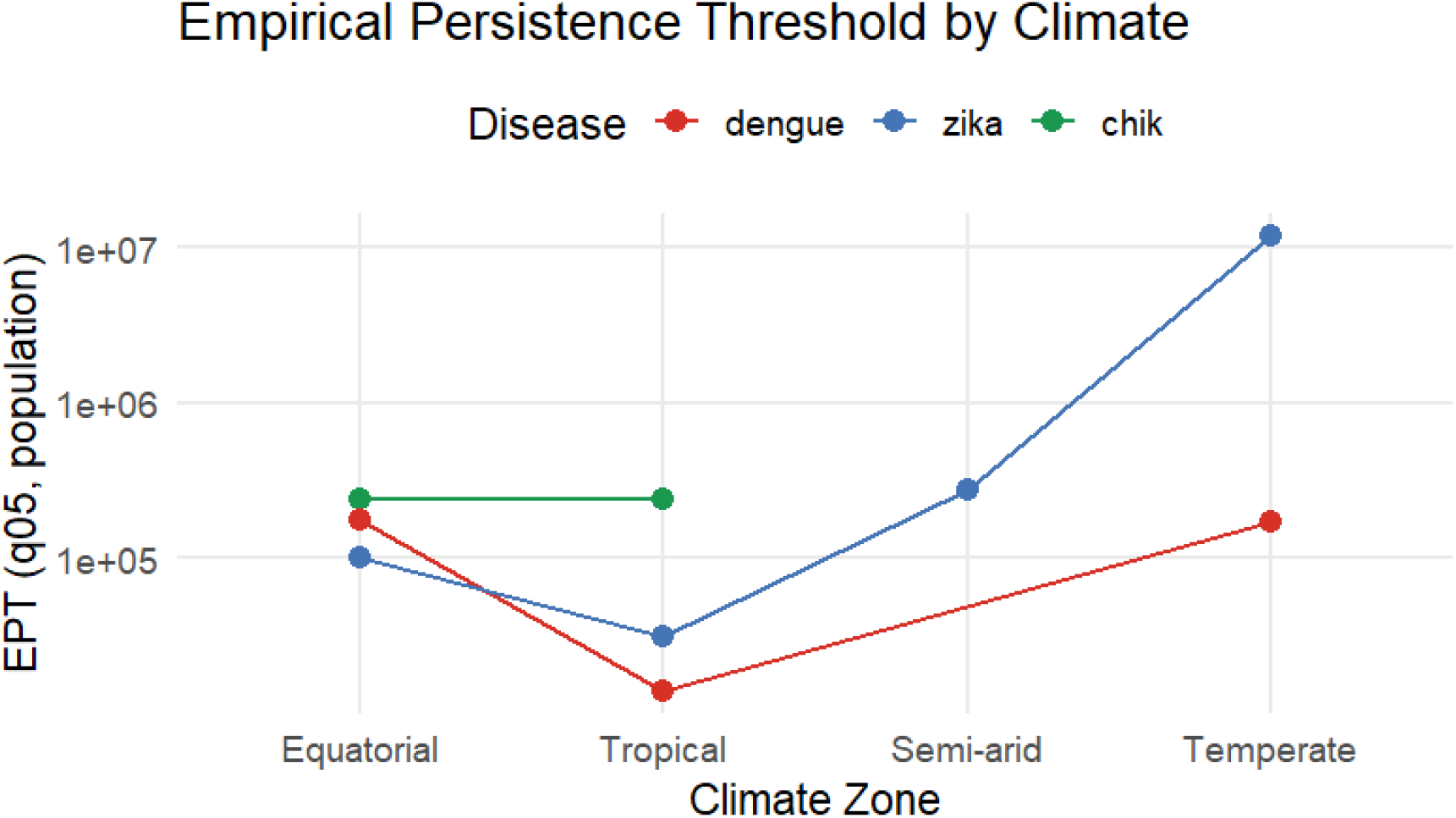
Empirical persistence thresholds (EPT, *q*_0.05_) across climate zones for dengue, Zika, and chikungunya. Values are shown on a logarithmic scale. Zika exhibits strong variation across climates, reaching markedly higher thresholds in temperate regions, whereas dengue and chikungunya show more moderate variation.

For dengue, lower EPT values were consistently observed in tropical and equatorial climates, where temperature and precipitation conditions favor vector abundance and year-round transmission. In these regions, even relatively small municipalities were able to sustain persistent transmission. In contrast, semi-arid and temperate regions exhibited higher EPT values, indicating that larger populations are required to compensate for less favorable environmental conditions. Zika showed a similar but less pronounced pattern, with moderate sensitivity to climatic conditions. Chikungunya, however, displayed comparatively weak variation across climate classes, suggesting that population size plays a more dominant role in determining persistence for this disease.

### 3.6 Role of urban hierarchy (REGIC)

Urban hierarchy, as captured by the REGIC classification, also played a significant role in shaping persistence patterns. For dengue, high-persistence municipalities were not restricted to the most central nodes of the urban network. Instead, persistent transmission was observed across a wide range of REGIC levels, including peripheral municipalities.

This indicates that dengue can sustain transmission even in less connected urban contexts.

In contrast, chikungunya persistence was strongly concentrated in higher-level urban centers. Municipalities classified as metropolises or regional capitals accounted for the majority of high-persistence cases, reinforcing the importance of population size and connectivity.

Zika exhibited intermediate behavior, with persistence occurring in both central and intermediate urban levels, but less frequently in peripheral municipalities.

Figure 6 shows the variation of empirical persistence thresholds across urban hierarchy levels.

**Figure 6:**
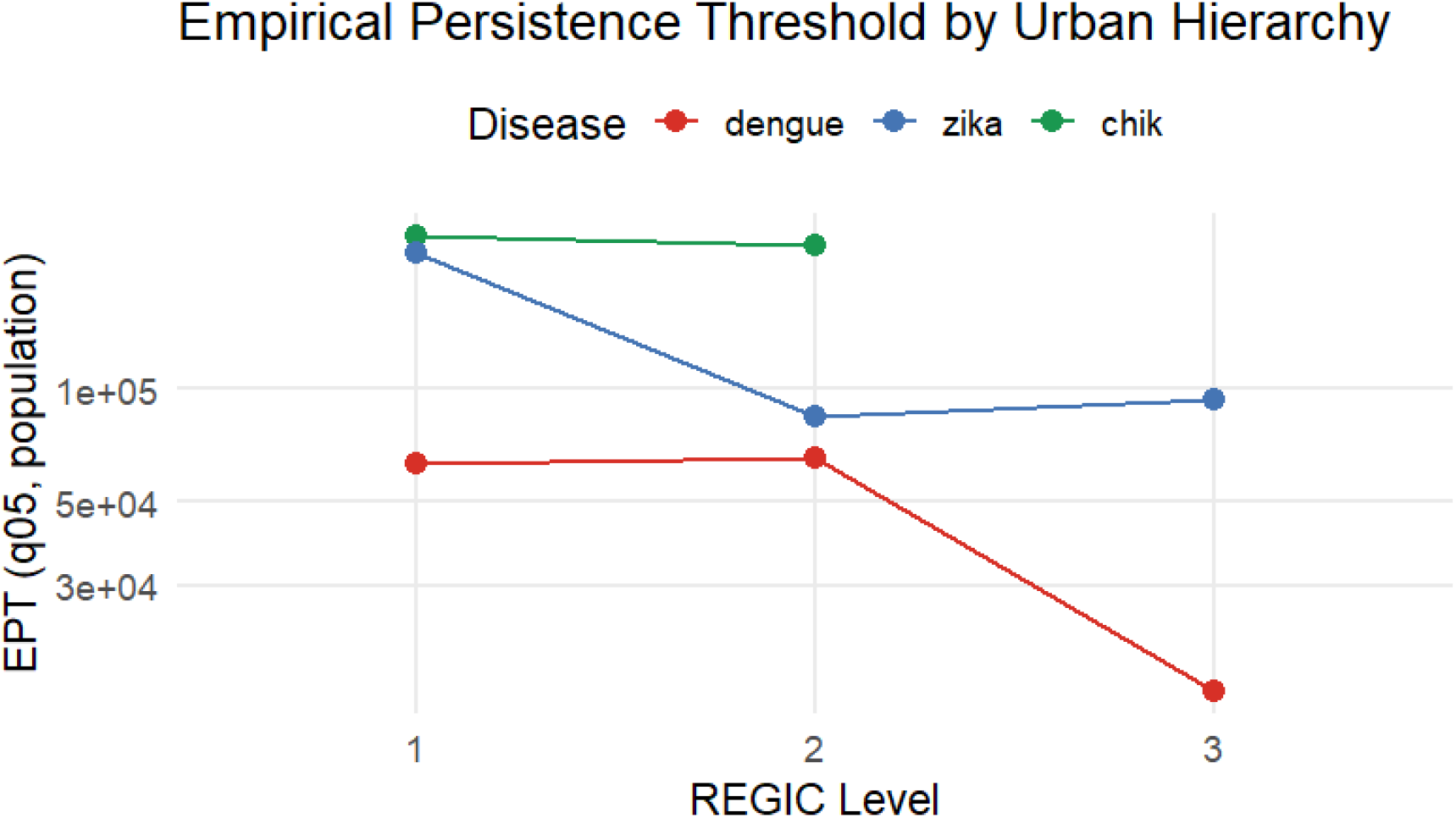
Empirical persistence thresholds (EPT, *q*_0.05_) across levels of the Brazilian urban hierarchy (REGIC) for dengue, Zika, and chikungunya. Values are shown on a logarithmic scale. Dengue and Zika exhibit non-monotonic patterns across hierarchical levels, whereas chikungunya remains consistently associated with higher thresholds.

Interestingly, large metropolitan areas did not always correspond to the highest persistence levels. Instead, they often exhibited greater variability, with transitions between persistence regimes. This suggests that high connectivity may simultaneously facilitate reintroduction and promote dynamical variability through the interplay between importation, susceptible depletion, and control interventions.

Detailed stratified results are provided in the supplementary material (S1 Table).

### 3.7 Synthesis: persistence as a multidimensional phenomenon

Across all analyses, a consistent picture emerges: epidemiological persistence in arboviral systems cannot be described by a single population threshold.

Instead, persistence reflects the interaction between multiple factors:

- population size, which sets the demographic scale of transmission;
- climate, which modulates vector dynamics and transmission efficiency;
- urban hierarchy, which captures connectivity within the broader metapopulation system.

The spatial patterns shown in Figure 7 reinforce the role of climate and urban connectivity in shaping persistence regimes.

**Figure 7:**
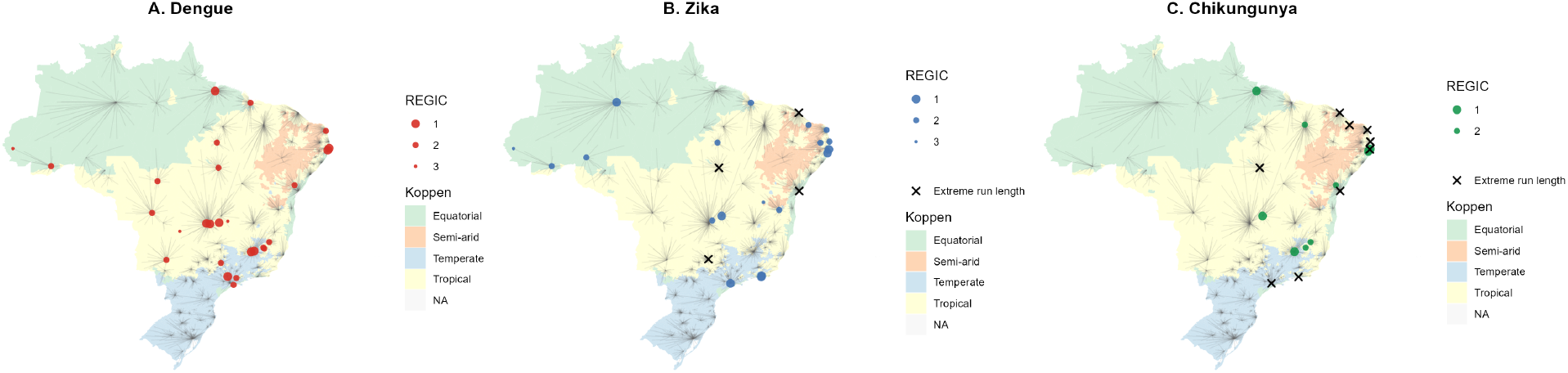
Spatial distribution of persistence regimes across Brazilian municipalities. Maps show municipalities classified according to persistence dynamics for dengue (A), Zika (B), and chikungunya (C). Colored circles indicate municipalities in the high-persistence regime, with symbol size proportional to the collapsed REGIC level. Black crosses denote municipalities with extreme run-length profiles. Background shading represents Köppen climate zones, and gray lines indicate connections in the urban network.

The Empirical Persistence Threshold provides a practical way to quantify this interaction, replacing the classical notion of a universal CCS with a probabilistic, data-driven description of persistence that varies across diseases and contexts.

## 4 Discussion

The classical notion of Critical Community Size (CCS) assumes that persistence can be described by a single population threshold separating endemic transmission from stochastic extinction. Our results show that this picture is no longer adequate for arboviruses circulating in highly connected urban systems.

Across all three diseases, persistence does not emerge as a sharp transition but rather as a structured continuum. The identification of two dominant regimes, low persistence and high persistence, provides a useful coarse-grained description, but the underlying dynamics are clearly more nuanced. Municipalities do not simply lie above or below a threshold; instead, they occupy positions along a gradient shaped by the temporal organization of transmission.

The methodological approach introduced here, based on run-length distributions and Wasserstein distances, allows this structure to be explicitly quantified. By focusing on the duration of consecutive transmission episodes, rather than on incidence magnitude, the analysis captures an aspect of epidemiological dynamics that is directly related to persistence. The use of the Wasserstein metric is particularly important in this context, as it preserves the temporal ordering of run lengths and provides a meaningful notion of distance between persistence profiles. This combination offers a simple but powerful framework for comparing transmission dynamics across large numbers of municipalities. Within this framework, the Empirical Persistence Threshold (EPT) emerges as a natural generalization of CCS. Instead of a single deterministic cutoff, EPT characterizes the lower tail of the population distribution among municipalities that sustain transmission. In this sense, it provides a probabilistic description of persistence that is more consistent with the stochastic and spatially coupled nature of modern epidemiological systems.

The results reveal clear and systematic differences across arboviruses. Dengue exhibits the lowest persistence thresholds and is able to sustain transmission even in relatively small municipalities. This is consistent with its longterm establishment in Brazil and its strong adaptation to a wide range of ecological and urban conditions. At the other extreme, chikungunya shows markedly higher thresholds, indicating that sustained transmission is largely restricted to larger population centers. Zika occupies an intermediate position, suggesting that its persistence dynamics are more sensitive to both demographic and environmental factors.

Climate plays an important but heterogeneous role. For dengue, favorable climatic conditions, particularly in tropical and equatorial regions, substantially lower the population scale required for persistence. In less favorable climates, larger pospulations are needed to maintain transmission. Zika shows a similar, though weaker, dependence. In contrast, chikungunya appears to be less sensitive to climatic variation, with persistence primarily driven by population size.

Urban hierarchy, as captured by the REGIC classification, adds an additional layer of structure. For dengue, persistent transmission is observed across a wide range of urban levels, including relatively peripheral municipalities. This suggests that dengue is capable of maintaining local transmission even in less connected settings. For chikungunya, persistence is much more concentrated in higher-level urban centers, reinforcing the importance of large, well-connected populations. Zika again shows intermediate behavior.

An interesting and somewhat counterintuitive result is that the largest metropolitan areas do not necessarily exhibit the most stable persistence. Instead, they often display higher variability and transitions between regimes. This suggests that strong connectivity may have a dual effect: it facilitates rapid reintroduction of the pathogen, but also contributes to dynamical instability through complex interactions between importation, depletion of susceptible individuals, and intervention measures.

Taken together, these findings point to a broader conclusion: persistence in modern urban systems is an emergent property resulting from the interaction between population size, environmental suitability, and network structure. No single variable is sufficient to explain the observed patterns. In particular, the idea of a universal critical population threshold is not supported by the data.

The EPT framework provides a practical way to operationalize this perspective. It allows persistence to be quantified in a manner that is both empirically grounded and directly comparable across diseases and contexts. At the same time, it remains simple enough to be applied to large surveillance datasets without requiring detailed mechanistic modeling.

Some limitations should be noted. The analysis is based on reported case data, which may be affected by underreporting and variation in surveillance quality across municipalities. The binary representation of incidence, while robust, does not capture differences in transmission intensity. In addition, the classification into persistence regimes depends on clustering choices, although the two-regime structure proved to be stable across analyses.

Despite these limitations, the results consistently support a reinterpretation of persistence as a probabilistic and context-dependent phenomenon. This has practical implications for public health. Interventions aimed at reducing transmission cannot rely solely on population size as an indicator of risk. Instead, they must account for environmental conditions and the position of municipalities within the urban network.

In conclusion, this study shows that pathogen persistence in highly connected urban systems cannot be adequately described by a single deterministic threshold. By combining run-length representations with optimal transport metrics, and by introducing the Empirical Persistence Threshold, we provide a framework that captures the multidimensional nature of persistence. This approach offers a bridge between classical epidemiological theory and the realities of contemporary, data-rich, spatially connected disease systems.

## Acknowledgments

The authors used generative artificial intelligence (ChatGPT) to assist with language refinement and manuscript organization. All conceptual development, data analysis, and scientific conclusions were independently performed and validated by the authors.

## Data Availability

All data used in this study are publicly available from the Brazilian Ministry of Health through the DATASUS platform (https://datasus.saude.gov.br). Processed datasets and scripts used for analysis are available from the corresponding author upon reasonable request.

## Ethics Statement

This study used publicly available aggregated data and did not involve human subjects. Therefore, ethical approval was not required.

